# Tissue-like structures formed by a bacterium

**DOI:** 10.1101/2025.06.27.661998

**Authors:** Gillian M. L. Ampah, Charles J. Myers, Carlos A. Ramírez Carbó, Isbella S. Lin, Beiyan Nan

## Abstract

Bacteria generally form only simple multicellular structures lacking the stable cell-cell connections characteristic of eukaryotic tissues. However, when the antibiotic moenomycin modifies peptidoglycan cell wall synthesis, rod-shaped cells of the Gram-negative bacterium *Myxococcus xanthus* become spherical, fuse their outer membranes, and assemble into stable, honeycomb-like lattices resembling eukaryotic tissues. These findings raise the intriguing possibility that some tissue-like organization could have evolved from stress-induced responses in bacterial ancestors.

---

The shift from single cells to multicellular tissues marks one of the most pivotal milestones in evolution. Multicellularity has arisen independently on at least 25 occasions across diverse evolutionary lineages, with particularly prominent examples in plants, animals, and fungi ^1^. Bacteria can also form multicellular structures, with myxobacteria standing out for their well-documented multicellular behavior. The model organism *Myxococcus xanthus* migrates in large, coordinated swarms across surfaces—a dynamic form of biofilm. Under starvation, these swarms give rise to fruiting bodies containing millions of spores encased in protective polysaccharide shells ^2^. However, bacterial multicellular structures differ fundamentally from eukaryotic tissues due to their lack of stable cell-cell connections. While relatively persistent assemblies like biofilms and fruiting bodies are maintained by extracellular matrices, they lack the direct intercellular contacts characteristic of true tissues ^3,4^. Conversely, direct connections—such as those formed through inner membrane (**IM**) nanotubes, outer membrane (**OM**) fusion, or various secretion systems—are typically transient and insufficient to support tissue-like architecture ^5-8^.

When PG synthesis in the rod-shaped *M. xanthus* (**Fig. 1a**)is stressed by the antibiotic moenomycin, the cells do not lyse; instead, they convert into spheres (**Fig. 1b**) ^9^. Unlike L-forms that lack the peptidoglycan (**PG**) cell wall, often depend on osmo-protectants for survival ^10^, and generate heterogenous, bubble-like progenies ^11^, these spheres can grow in low-osmolarity media and are uniform in size ^9^. Notably, they continued to synthesize PG, as demonstrated by the incorporation of TAMRA 3-amino-D-alanine (**TADA**), a fluorescent D-amino acid, by the enzymes that assemble the PG matrix (**Fig. 2**) ^12^.

**Fig. 1.**
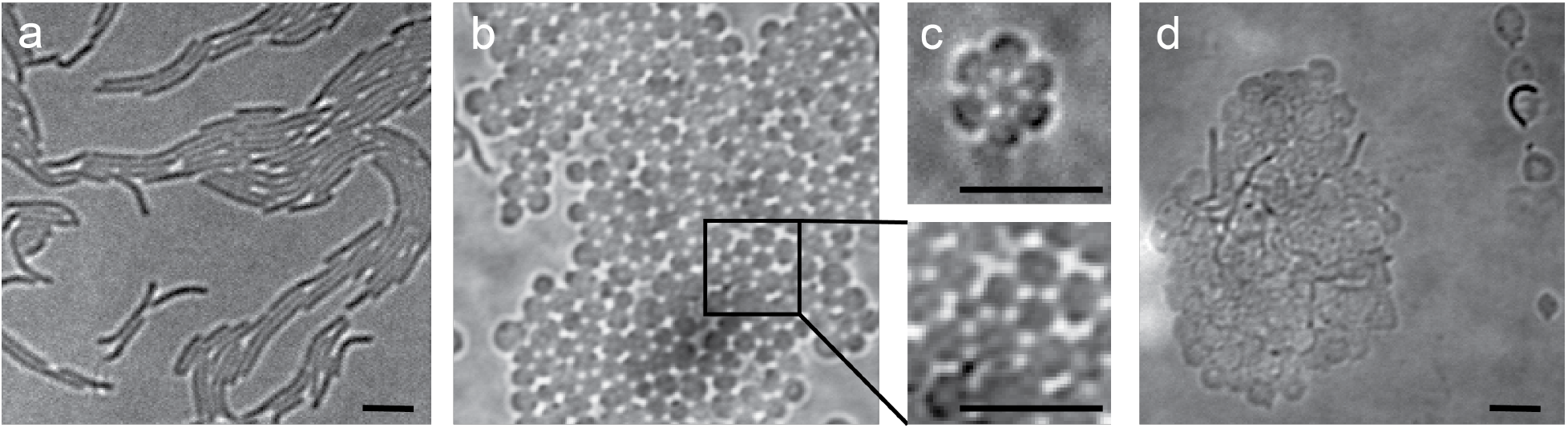
*M. xanthus* cells form tissue-like structures under moenomycin stress. (**a**) Wild-type vegetative cells are rod-shaped. (**b, c**) After moenomycin treatment (4 μg/ml, 4 h), cells transform into spheres and form honeycomb-like lattices of different sizes through visible cell-cell connections. (**d**) Cells overexpressing Pal lysed frequently and did not form tissue-like structures after moenomycin treatment. Scale bars, 5 μm.

**Fig. 2.**
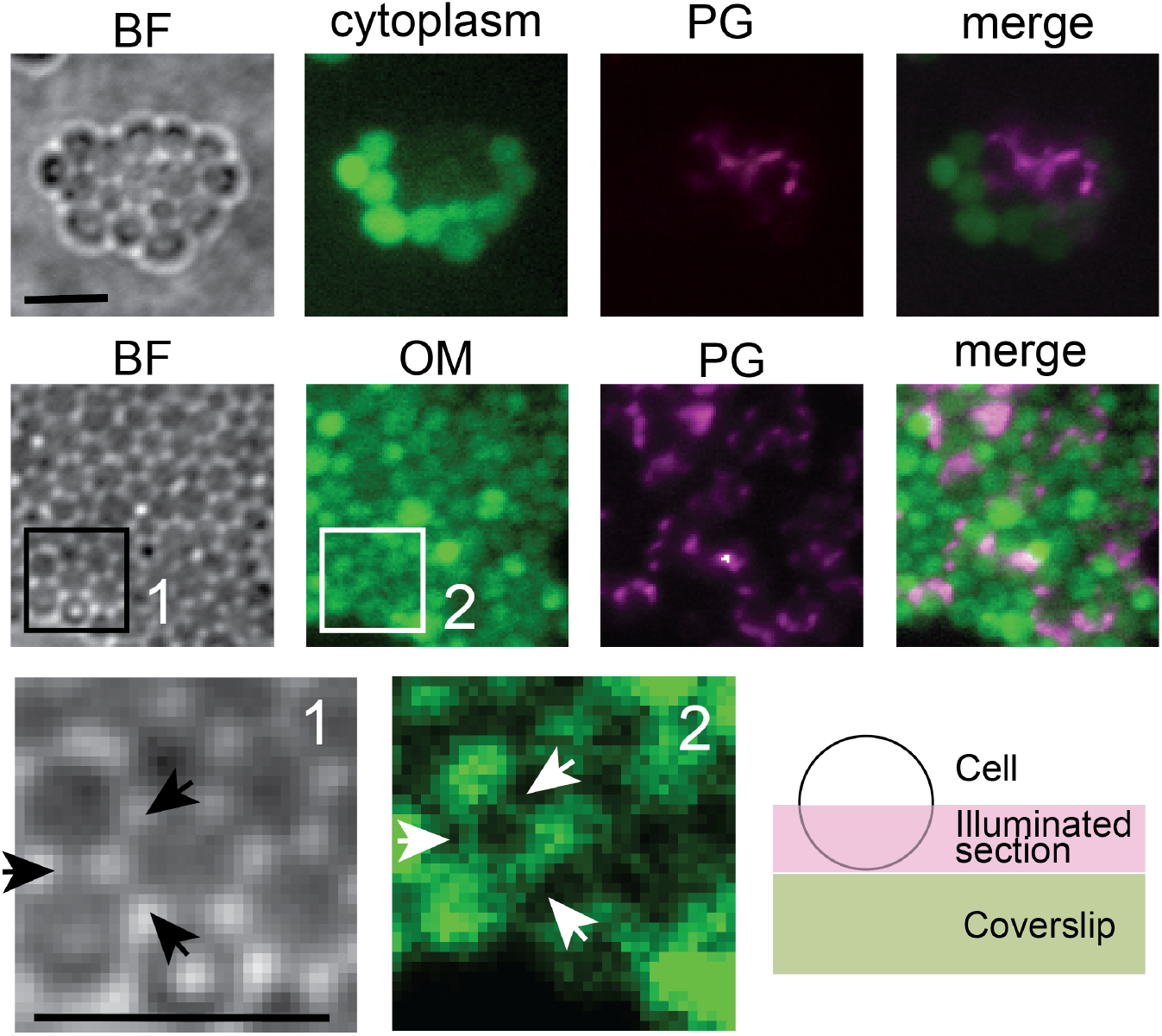
Cells in the tissue-like structures connect to each other through extensive OM fusion. Cells share their OMs but not cytoplasms. The cytoplasm, PG, and OM were labeled with calcein, TADA, and WGA, respectively. BF, bright field. Arrows point to the intracellular OM connections. Scale bars, 5 μm. The cartoon inset depicts the cell sections imaged using HILO illumination.

Remarkably, these spheres organized into lattices of varying sizes on agar surfaces, displaying a hexagonal cellular arrangement with bridge-like connections linking adjacent cells (**Fig. 1, 2**). These lattices, reminiscent of eukaryotic tissues, exhibited sustained growth over time on agar surfaces. To determine whether individual cells within the lattices retained their integrity, we treated a liquid culture of wild-type *M. xanthus* with moenomycin and split it into two samples: one incubated with calcein and the other with TADA. Calcein is a small, nonfluorescent, hydrophobic molecule that readily crosses IMs. It serves as an effective probe for IM integrity, as cytoplasmic esterases convert it into a fluorescent, hydrophilic product that becomes trapped inside the cell, accumulating in the cytoplasm ^6^. After 1 h of incubation, samples were washed, mixed at a 1:1 ratio and spotted on an agar surface.

Each sample incorporated its respective dye in liquid culture, and upon mixing, they quickly assembled into honeycomb-like lattices on the agar surface. We used the highly inclined and laminated optical sheet (HILO) illumination ^13,14^ to image half of the cell surface that is close to the coverslips. After one hour of incubation, no calcein leakage was observed, indicating that the cells remained viable and preserved IM integrity. Importantly, TADA-labeled cells showed no fluorescence from calcein, suggesting that cytoplasmic contents were not exchanged between individual cells (**Fig. 2**).

*M. xanthus* is a typical Gram-negative bacterium characterized by the presence of an OM surrounding its PG layer. To assess whether cells could establish connections via their OMs, we conducted a similar tissue-forming experiment using cells labeled with different markers. In this setup, calcein was replaced with Alexa Fluor 488-conjugated wheat germ agglutinin (**WGA**), which cannot penetrate the OM and therefore selectively stains OM ^15^. In contrast to the lack of cytoplasmic exchange, TADA-labeled cells rapidly acquired fluorescence from WGA, resulting in its nearly uniform distribution in the tissue-like lattices (**Fig. 2**). Consistent with our recent discovery that the components in *M. xanthus* OM diffuse freely ^16^, this result indicate that individual cells share OMs in the tissue-like structures. Strikingly, the “bridges” between cells were also stained by WGA (**Fig. 2**), indicating that the tissue-like structures are organized by stable OM channels.

In untreated swarms, *M. xanthus* cells can transiently fuse their OMs in a limited scale ^17^. But how does moenomycin—an antibiotic that modulates PG synthesis ^18^— promote the formation of a stable and extensive OM network? Given that Gram-negative bacteria typically anchor their OMs to the PG layer to coordinate cell wall elongation and division, we hypothesize that moenomycin-induced alterations to PG structure weaken these tethering points, thereby enabling extensive OM fusion. To test this hypothesis, we sought to reinforce the PG-OM connection by overexpressing Pal—a protein that binds peptidoglycan noncovalently and links it to outer membrane components of the Tol-Pal system ^19^—using a vanillate-inducible promoter activated with 100 µM sodium vanillate. Following moenomycin treatment, cells still adopted a spherical morphology but were unable to form tissue-like structures (**Fig. 1d**), indicating that reinforced PG-OM connections inhibit tissue formation. Notably, widespread cell lysis was observed (**Fig. 1d**). Given that soil, its natural habitat, is abundant in antimicrobials produced by *M. xanthus* itself and other organisms like actinobacteria, forming tissue-like structures is likely a survival mechanism.

It is often assumed that tissue formation arose through gain-of-function events during evolution. However, our findings suggest a counterintuitive possibility: the emergence of primitive tissue-like structures may have resulted from loss-of-function events in bacterial cell wall assembly. Bacteria have long been implicated in key evolutionary transitions toward multicellularity, with the origin of animal tissues being particularly significant due to its relevance to human biology. For example, *Salpingoeca rosetta*, a unicellular eukaryote closely related to animals, forms rosette-shaped colonies in response to specific prey bacteria ^20^. Myxobacteria— the apex predatory microbes with complex behaviors—exhibit foraging strategies reminiscent of early animals ^2^ and uniquely produce plasmalogens, lipids otherwise exclusive to animals ^21^. Phylogenetic evidence also suggests that ancient myxobacteria contributed to the development of the first eukaryotic cytoplasm ^22^. These insights raise the intriguing possibility that animal tissues may have evolved from stress-induced responses in bacterial ancestors.

## Materials and methods

Vegetative *M. xanthus* cells were grown in liquid CYE medium (10 mM MOPS pH 7.6, 1% (w/v) Bacto™ casitone (BD Biosciences), 0.5% yeast extract and 8 mM MgSO^4^) at 32 °C, in 125-ml flasks with vigorous shaking, or on CYE plates that contains 1.5% agar. We used strain DZ2 as the wild-type *M. xanthus* strain. Pal Overexpression strain was constructed by electroporating DZ2 cells with 4 µg of plasmid DNA. The *pal* gene, driven by a vanillate-inducible promoter, was inserted into the Mx4 phage attachment site as a merodiploid on the *M. xanthus* chromosome. Transformed cells were plated on CYE plates supplemented with 10 mg/ml tetracycline and expression was induced by 100 µM sodium vanillate. For microscopy imaging, cultures were grown in liquid CYE to OD^600^ ∼1 and supplemented with 10 µg/mL WGA Alexa Fluor-488 conjugate and 75 µM TADA for one hour. Cells were spun down at 6,000x g for 3 min and the pellet washed three times with CYE. 5 µl of cells were spotted on agar (1.5%) pads and imaged using a Andor iXon Ultra 897™ EMCCD camera (effective pixel size 160 nm) on an inverted Nikon Eclipse-Ti™ microscope with a 100× 1.49 NA TIRF objective. Fluorescence of TADA and Alexa Fluor 488-conjugated WGA was excited by the 561-nm and 488-nm lasers, respectively.

## Acknowledgments

We thank Drs. Michael Van Nieuwenhze and Yen-Pang Hsu for providing TADA and all the members in the Nan laboratory for technical support and constructive discussions. Part of this work was supported by the National Institutes of Health grants GM129000 to B. N..

## Notes

### Competing Interest Statement

The authors have declared no competing interest.

## References

1 Grosberg, R. K. & Strathmann, R. R. The evolution of multicellularity: A minor major transition? Annu Rev Ecol Evol S 38, 621–654 (2007). 10.1146/annurev.ecolsys.36.102403.114735

2 Kroos, L. et al. Milestones in the development of Myxococcus xanthus as a model multicellular bacterium. Journal of Bacteriology 0, e00071–00025 10.1128/jb.00071-25

3 O’Connor, K. A. & Zusman, D. R. Reexamination of the role of autolysis in the development of Myxococcus xanthus. J Bacteriol 170, 4103–4112 (1988).

4 Flemming, H. C. & Wingender, J. The biofilm matrix. Nat Rev Microbiol 8, 623–633 (2010). 10.1038/nrmicro2415

5 Angulo-Canovas, E. et al. Direct interaction between marine cyanobacteria mediated by nanotubes. Sci Adv 10, eadj1539 (2024). 10.1126/sciadv.adj1539

6 Dubey, G. P. & Ben-Yehuda, S. Intercellular nanotubes mediate bacterial communication. Cell 144, 590–600 (2011). 10.1016/j.cell.2011.01.015

7 Vassallo, C. N. et al. Infectious polymorphic toxins delivered by outer membrane exchange discriminate kin in myxobacteria. Elife 6 (2017). 10.7554/eLife.29397

8 Basler, M., Ho, B. T. & Mekalanos, J. J. Tit-for-tat: type VI secretion system counterattack during bacterial cell-cell interactions. Cell 152, 884–894 (2013). 10.1016/j.cell.2013.01.042

9 Zhang, H., Venkatesan, S., Ng, E. & Nan, B. Coordinated peptidoglycan synthases and hydrolases stabilize the bacterial cell wall. Nature communications 14, 5357 (2023). 10.1038/s41467-023-41082-3

10 Errington, J., Mickiewicz, K., Kawai, Y. & Wu, L. J. L-form bacteria, chronic diseases and the origins of life. Philos Trans R Soc Lond B Biol Sci 371 (2016). 10.1098/rstb.2015.0494

11 Mercier, R., Kawai, Y. & Errington, J. Wall proficient E. coli capable of sustained growth in the absence of the Z-ring division machine. Nat Microbiol 1, 16091 (2016). 10.1038/nmicrobiol.2016.91

12 Hsu, Y. P. et al. Full color palette of fluorescent d-amino acids for in situ labeling of bacterial cell walls. Chem Sci 8, 6313–6321 (2017). 10.1039/c7sc01800b

13 Fu, G. et al. MotAB-like machinery drives the movement of MreB filaments during bacterial gliding motility. Proc Natl Acad Sci U S A 115, 2484–2489 (2018). 10.1073/pnas.1716441115

14 Tokunaga, M., Imamoto, N. & Sakata-Sogawa, K. Highly inclined thin illumination enables clear single-molecule imaging in cells. Nat Methods 5, 159–161 (2008). 10.1038/nmeth1171

15 Dubois, L., Vettiger, A., Buss, J. A. & Bernhardt, T. G. Using fluorescently labeled wheat germ agglutinin to track lipopolysaccharide transport to the outer membrane in Escherichia coli. mBio 16, e0395024 (2025). 10.1128/mbio.03950-24

16 Topo, E. J., De Santiago, C. B., Cao, P., Wall, D. & Nan, B. Mechanism of bacterial outer membrane exchange revealed by quantitative microscopy. bioRxiv, 2025.2004.2025.650704 (2025). 10.1101/2025.04.25.650704

17 Cao, P. & Wall, D. Direct visualization of a molecular handshake that governs kin recognition and tissue formation in myxobacteria. Nature communications 10, 3073 (2019). 10.1038/s41467-019-11108-w

18 Lovering, A. L., de Castro, L. H., Lim, D. & Strynadka, N. C. Structural insight into the transglycosylation step of bacterial cell-wall biosynthesis. Science 315, 1402–1405 (2007). 10.1126/science.1136611

19 Szczepaniak, J. et al. The lipoprotein Pal stabilises the bacterial outer membrane during constriction by a mobilisation-and-capture mechanism. Nature communications 11, 1305 (2020). 10.1038/s41467-020-15083-5

20 Alegado, R. A. et al. A bacterial sulfonolipid triggers multicellular development in the closest living relatives of animals. Elife 1, e00013 (2012). 10.7554/eLife.00013

21 Gallego-Garcia, A. et al. A bacterial light response reveals an orphan desaturase for human plasmalogen synthesis. Science 366, 128–132 (2019). 10.1126/science.aay1436

22 Lopez-Garcia, P. & Moreira, D. The Syntrophy hypothesis for the origin of eukaryotes revisited. Nat Microbiol 5, 655–667 (2020). 10.1038/s41564-020-0710-4

